# Olfactory system structure and function in newly hatched and adult locusts

**DOI:** 10.1101/2023.11.27.568838

**Authors:** Kui Sun, Subhasis Ray, Nitin Gupta, Zane Aldworth, Mark Stopfer

## Abstract

An important question in neuroscience is how sensory systems change as animals grow and interact with the environment. Exploring sensory systems in animals as they develop can reveal how networks of neurons process information as the neurons themselves grow and the needs of the animal change. Here we compared the structure and function of peripheral parts of the olfactory pathway in newly hatched and adult locusts. We found that populations of olfactory sensory neurons (OSNs) in hatchlings and adults responded with similar tunings to a panel of odors. The morphologies of local neurons (LNs) and projection neurons (PNs) in the antennal lobes (ALs) were very similar in both age groups, though they were smaller in hatchlings, they were proportional to overall brain size. The odor evoked responses of LNs and PNs were also very similar in both age groups, characterized by complex patterns of activity including oscillatory synchronization. Notably, in hatchlings, spontaneous and odor-evoked firing rates of PNs were lower, and LFP oscillations were lower in frequency, than in the adult. Hatchlings have smaller antennae with fewer OSNs; removing antennal segments from adults also reduced LFP oscillation frequency. Thus, consistent with earlier computational models, the developmental increase in frequency is due to increasing intensity of input to the oscillation circuitry. Overall, our results show that locusts hatch with a fully formed olfactory system that structurally and functionally matches that of the adult, despite its small size and lack of prior experience with olfactory stimuli.

## Introduction

The sense of smell, processed by the olfactory system, is essential for survival throughout the lifespan of most animals because it plays critical roles in behaviors such as foraging, avoiding predators, and finding mates (Fischer et al., 2017; Togunov et al., 2017; Zjacic & Scholz, 2022). During development an animal’s relationship with the olfactory environment can profoundly transform as sources of, and preferences for, food frequently change (Nakazawa, 2015), predation dangers can increase or decrease (Cooper, Jr & Stankowich, 2010; Ohlberger et al., 2019), and the onset of sexual maturity is often accompanied by increased sensitivity to pheromone cues involved in locating mates and courtship (Gadenne et al., 1993). How the olfactory system is modified as an animal develops can reveal fundamental aspects of information processing by networks of neurons. The study of sensory processing in relatively simple animals such as insects has proved to be a useful strategy for answering basic questions about these phenomena (Laurent & Davidowitz, 1994; Laurent, 1999; Laurent et al., 2001; MacLeod & Laurent, 1996; Mazor & Laurent, 2005; Perez-Orive et al., 2002; Stopfer et al., 1997; Stopfer & Laurent, 1999; Stopfer et al., 2003).

Flies, moths, and other holometabolous insects undergo dramatic anatomical and physiological changes as they undergo metamorphosis from the larval to the adult stage (Truman, 2019). In these animals, the diet and behavior of the larva can differ greatly from the adult (Truman & Riddiford, 1999). These changes are reflected in the development of their nervous system (Levine & Truman, 1982; Tissot & Stocker, 2000). For example, in the moth *Manduca sexta*, olfactory sensory neurons (OSNs) gradually begin to generate spontaneous and odor evoked activity (Boeckh & Tolbert, 1993; Schweitzer et al., 1976), and glomeruli in the antennal lobes (ALs) start to form only in the 6th stage of development. In contrast, hemimetabolous insects like locusts and cockroaches hatch with bodies in many ways resembling the adult and later undergo incomplete metamorphosis (Truman, 2019). Locust hatchlings, like adults, feed mainly on grass or other plants. Since locust hatchlings receive no parental care and may emerge at some distance from food sources, to survive, they need to forage independently.

As locusts grow from hatchling to adulthood, they acquire experience with the olfactory environment, their body size increases by an order of magnitude (Capinera & Squitier, 1996), and their brain size doubles. We have shown recently that newly hatched locusts lacking any prior exposure to food are innately attracted by food odors (Ray et al., 2023). Notably, earlier work in a related species, *Schistocerca gregaria*, established that newly hatched locusts have adult-like glomeruli in their ALs, and their OSNs and PNs respond by generating action potentials when odors are presented to the antennae (Anton et al., 2002).

Here we provide an analysis of the structure and function of the first stages in the olfactory pathway in newly hatched and adult *Schistocerca americana*. Our quantitative results indicate that hatchling olfactory OSNs, local and projection neurons (LNs and PNs) function like those of adults: they respond with similar tunings to panels of odors with complex, coordinated, oscillatory activity patterns. Although smaller, LNs and PNs in hatchlings resemble those in adults. Notably, odor-elicited neural oscillations gradually increase in frequency as the locusts age, a change we found can be explained by increased excitatory drive as new OSNs are added to growing antennae. Overall, our results show that locusts hatch with a fully formed olfactory system that structurally and functionally matches that of the adult, despite its small size and lack of prior experience with olfactory stimuli.

## Materials and Methods

### Animal subjects

The locust *Schistocerca americana* used in this study were reared in crowded colonies in our indoor lab. Mature females laid eggs inside cups filled with clean sand which were then incubated at 28.8 °C until the eggs hatched. We used locusts that were freshly hatched (1^st^ instar), 10 days old (3^rd^ instar), 20 days old (5^th^ instar), and adult locusts of either sex in our experiments.

### Odor stimuli

We tested a large panel of odorants including some associated with food sources and others that our locusts, raised in the laboratory, hadn’t encountered before. Odorants were mixed in mineral oil and delivered by an olfactometer as square pulses within a constant stream of filtered and humidified air to the intact antennae (Gupta & Stopfer, 2012). The odor panel included 1-hexanol (hex), cis-3-hexen-1-ol (cis), 1-hexanal (hxa), 1-octen-3-ol (oct), cyclohexanone (cyc), furfuryl mercaptan (fur), ethyl mercaptan (eth), citral (cit), pentyl acetate (pen), camphor (camp), and wheat grass juice (wgj). The concentrations of the odorants were adjusted in mineral oil to a partial vapor pressure of 0.024 (that of 1% hex); this procedure ensured similar numbers of molecules of each odorant were delivered. Wheat grass juice, which is a mixture of many volatiles, was not adjusted by vapor pressure. Grass juice was extracted from wheat grass (*Triticum aestivum*) grown in our laboratory as described (Ray et al., 2023). All pure odorants were obtained from Sigma-Aldrich.

### Electrophysiology

Electroantennograms (EAG) were obtained by inserting a saline-filled sharp glass electrode (resistance ∼10 MΩ) into the antenna and a silver-chloride ground electrode into one eye and were recorded by a DC amplifier (Model 440; Brown-Lee Precision, San Jose, CA). To make LFP and patch clamp recordings, a wax cup was built around the head to hold a bath of locust ringer solution, and a small section of the cuticle was then removed to expose the brain (Gupta & Stopfer, 2012). To record LFPs, a blunt glass, saline-filled electrode was placed above the calyx of the mushroom body. For whole-cell patch clamp recordings, patch electrodes were pulled from borosilicate glass capillaries by a pipette puller (Model P-97; Sutter Instruments), with the program parameters tuned to produce an electrode resistance around 6 MΩ, filled with locust internal solution (Laurent et al., 1993) that was adjusted to osmolarity of about 350. Data were recorded through a MultiClamp 700A microelectrode amplifier and digitized via Digidata 1322A. Stimulus delivery and recordings were controlled by our custom LabView program running on a data acquisition PC.

### Histology and imaging

In many experiments, either neurobiotin (Vector SP-1120) or lucifer yellow (Molecular Probes L453) was injected into recorded neurons at the conclusion of a session. Afterwards the brain was removed, fixed in 4% paraformaldehyde overnight, and then washed in phosphate buffered saline (PBS). In the case of neurobiotin, the brain was transferred into PBS with 3% Triton X and kept for an hour to permeabilize the membrane, followed by conjugation of neurobiotin with streptavidin Alexa Fluor 488, 568, or 633 (Invitrogen S11223, S11226, or S21375, respectively). Because lucifer yellow is itself fluorescent, brains labeled with it did not require the conjugation step. All brains were then dehydrated by an ethanol series, cleared with methyl salicylate and imaged under a confocal microscope. Neuronal morphologies were traced from the confocal image stacks using NeuroLucida, Simple Neurite Tracer plugin in imagej, or neutube software.

### Data analysis

Data analysis was carried out with custom written scripts in Matlab and Python. Signal processing and statistical tests were carried out using the functions from scipy and statsmodels packages in Python. All analysis scripts have been made available online.

Adult PNs have been shown to respond to odors with complex sequences of excitation and inhibition. As a measure of response complexity, we compared spiking epoch transitions in adult and young locusts. Spike rates in 400 ms wide bins were computed by simple histogram. A PN was considered responsive in a given bin if its firing rate within 4 s from odor onset differed from the background rate averaged across trials by more than one standard deviation. PN-odor pairs in which at least half the trials showed responses to odor puffs in the same bin were included for comparison of firing pattern complexity. We computed the firing rates in all 6 trials of these PN-odor pairs as the inverse of the inter-spike interval (ISI) and smoothed them with a Gaussian window with 33.33 ms standard deviation and 200 ms (6 standard deviations) width. A peak detection algorithm implemented in Python (https://gist.github.com/sixtenbe/1178136) was then used to identify local peaks in the firing rates. The peaks indicated excitatory epochs and the valleys between them indicated inhibitory epochs. Each valley counted for two transitions (excitatory-to-inhibitory and inhibitory-to-excitatory). If a peak appeared more than 200 ms after odor onset (or before the end of the response window, 4 s from odor onset), a spontaneous-to-excitatory (or excitatory-to-baseline) phase transition was counted. The distribution of transitions was compared between adult and hatchling locusts using Kolmogorov-Smirnov (KS) test, and showed no significant difference (KS=0.04, p=0.84).

To analyze LFP oscillations, signals were digitally bandpass filtered between 3-50 Hz. Spectrograms of the filtered signals were computed using a 0.5 s wide Hann window with 50% overlap between successive window positions. As successive segments of the antenna were removed, the spectrograms for trials 3-10 were averaged, and the frequency with the peak power within 1 s after odor onset was identified in this average spectrogram.

## Results

### Peripheral responses to odors are similarly tuned in hatchlings and adults

Antennae, the primary olfactory organs in insects, house OSNs in three different types of olfactory sensilla: basiconic, trichoidic, and coeloconic (Chapman, 2002; Ochieng et al., 1998). These OSNs initiate the olfaction process by generating spiking responses when odors enter their respective sensilla and bind to receptor molecules (Kim et al., 2023). We characterized the tuning of the OSN population in hatchlings and adults to a panel of odorants by recording electroantennograms (EAGs), which assess the summed responses of OSNs across the antenna. We found that EAGs from naïve hatchlings and adults displayed similar patterns of tuning (Figure 1). For example, hexanol, a prominent volatile released by wheat grass juice (Eissa et al., 2017; Hamilton-Kemp & Andersen, 1984; Njagi & Torto, 1996), evoked strong EAG responses in both adults and hatchlings, as did cis-3-hexen-1-ol, another wheat grass volatile known to evoke strong EAG response in other insects (Guerin & Visser, 1980; Njagi & Torto, 1996). In contrast, we observed much weaker EAG signals in both hatchlings and adults to pentanol, octen-3-ol, citral, and cyclohexanone. Pairwise t-tests with multiple-comparison correction showed no significant differences in response to any given odor between the two age groups, and a linear mixed model fit, a statistic appropriate for analyzing samples of unequal size, confirmed this result. Pairwise p-value for t-tests: cyc 0.86, wgj 0.08, cit 0.02, oct 0.75, pen 0.37, hxa 0.90, cis 0.34, where alpha was 0.006 after Bonferroni’s correction for multiple comparison (9 animals in each age-group, peak amplitude for hex, the normalizing value, was not compared; Table 1). For an ordinary least squares fit for the peak EAG amplitude as a function of age and stimulus using the formula “*amplitude = β0 + β1 x age + β2 x stimulus”*, where both age and stimulus were categorical variables, the coefficient *β1* for age was 0.004, and not significantly different from zero (p = 0.87, n=135 observations). Thus, we found that OSNs in the hatchling antennae can respond to a panel of odors with similar relative intensity profiles as those in the adult. This result suggests that age or experience with the olfactory environment does not substantially change the tuning of this population of neurons at the sensory periphery to general, non-pheromonal odorants.

**Figure 1:**
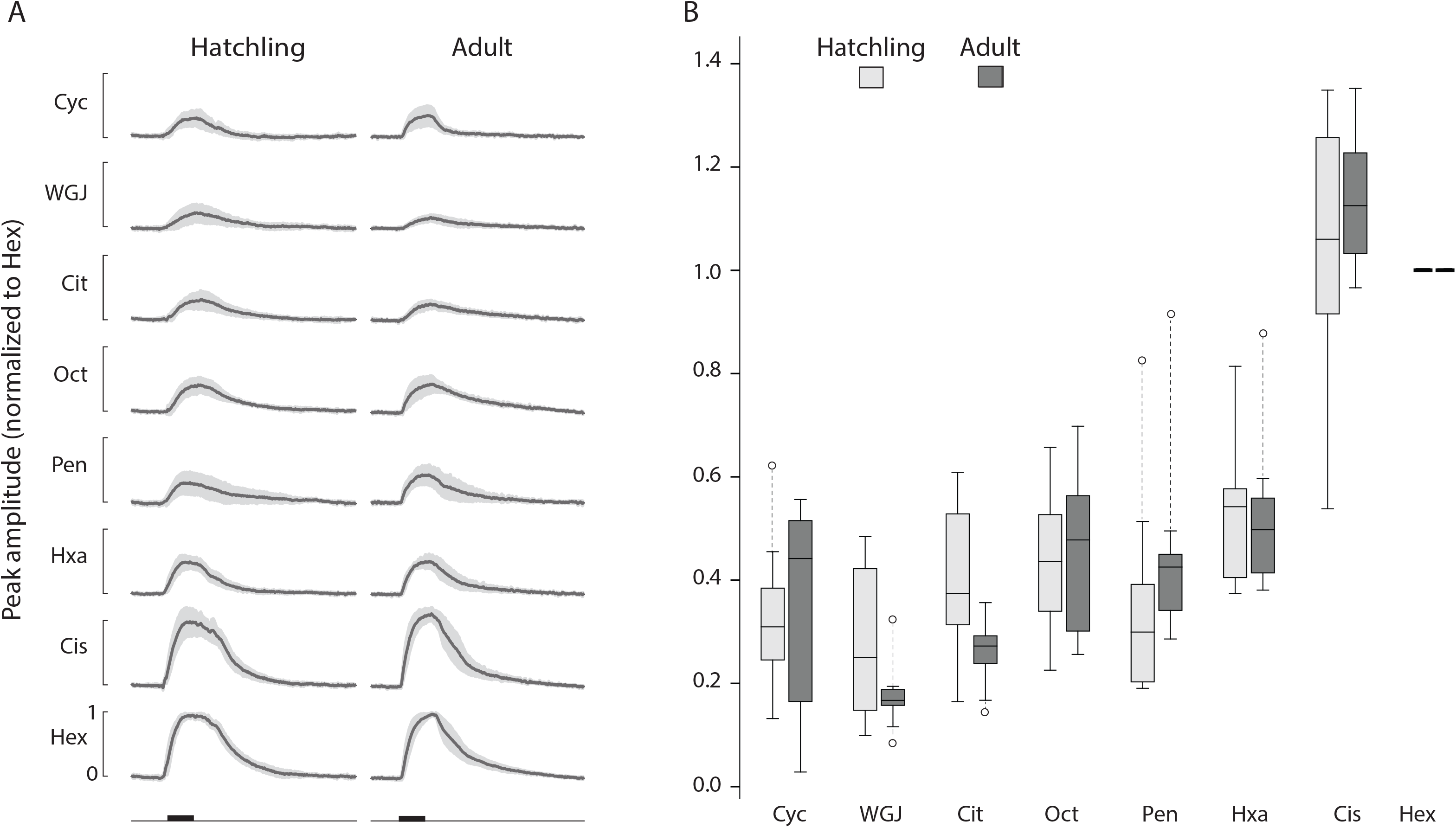
EAGs in adult (n=9) and freshly hatched locusts (n=8) show the ORN populations in hatchling and adults respond similarly to odors. (a) Average EAG responses to a panel of odors normalized by the maximum deflection elicited by hexanol in the same animal (black line). Gray band: 95% confidence level. Black horizontal bar: 1 sec odor pulse. (b) Distribution of normalized peak EAG amplitude for each odor across animals. Horizontal lines in the middle: median; whiskers: 1.5 interquartile range; circles: values outside this range.

**Figure 2:**
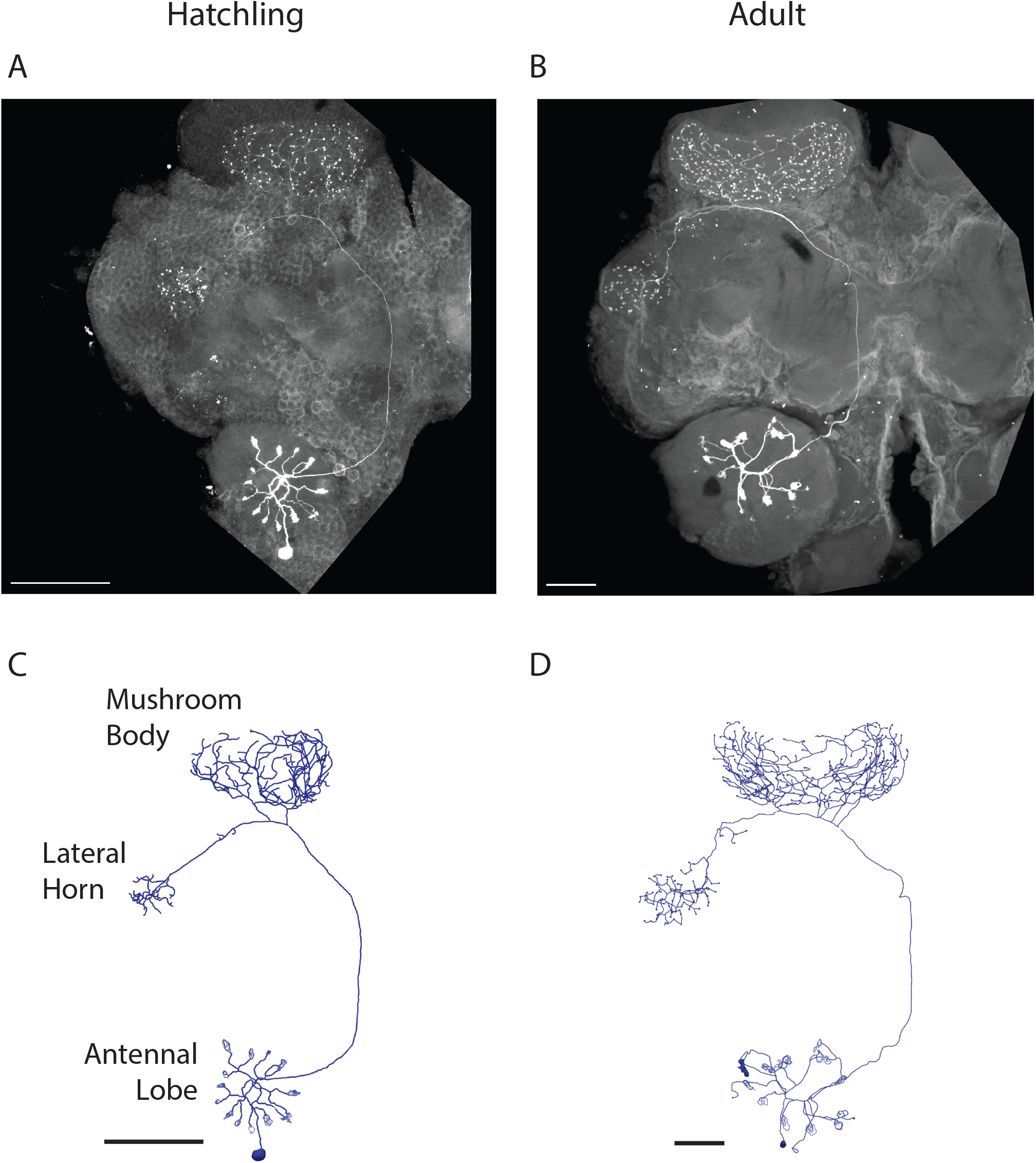
Morphologies of PNs in the locust AL are similar between hatchlings and adults. Shown are representative examples; see Table 2 for detailed comparisons. (a) Maximum intensity projections of hatchling and (b) adult hemibrains with dye-filled PNs. (c) Traced and reconstructed morphologies of the same hatchling and (d) adult PNs. Branches in the MB calyx are colored red. Blue dots indicate spines. Scale bar: 100 μm.

**Table 1:**
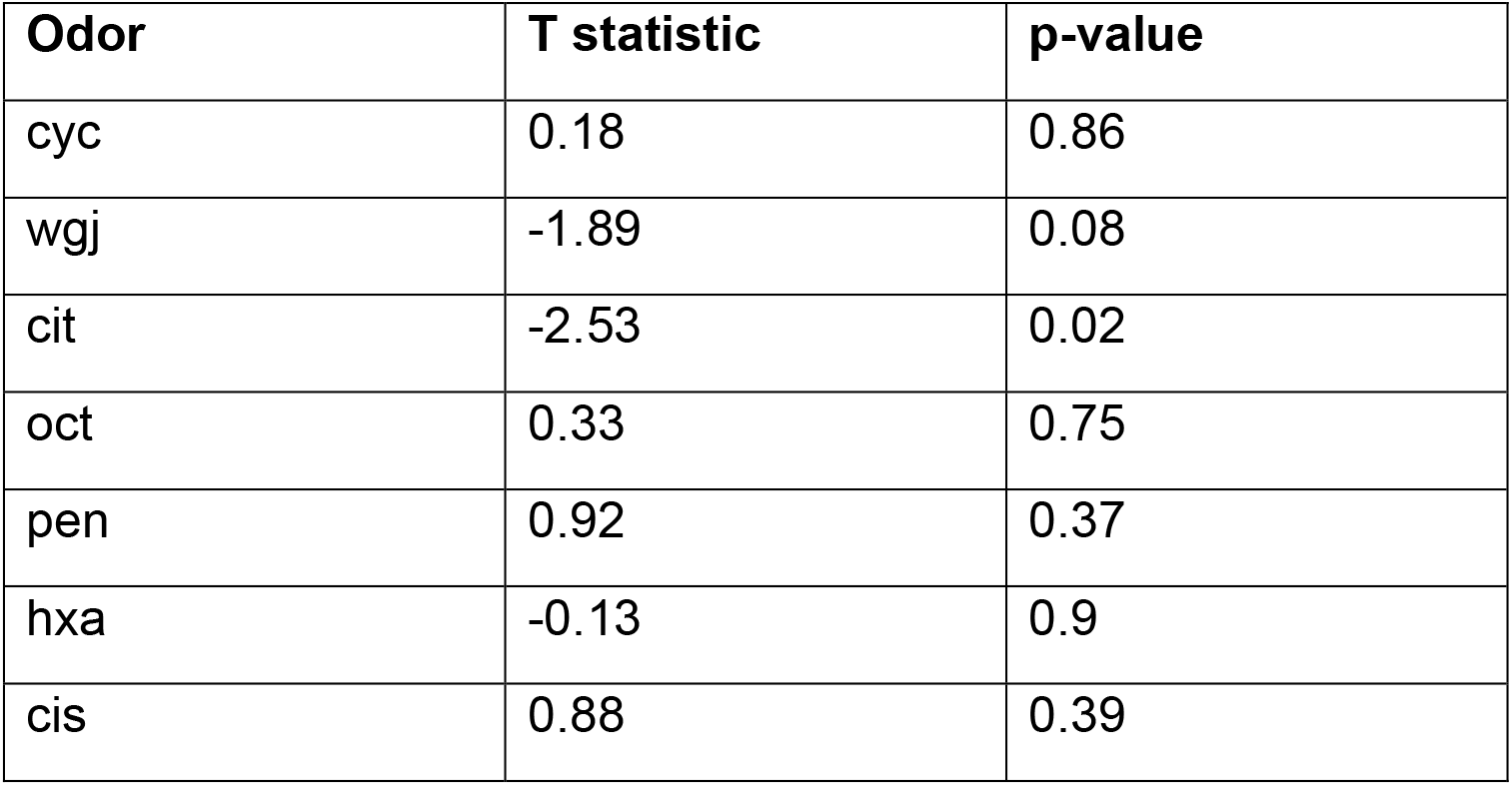
Quantitative comparison of EAG response peaks for various odors in freshly hatched and adult. All odors had 9 samples from each age group.

### Hatchling and adult AL neurons have similar morphologies

To compare olfactory antennal lobe neurons in hatchlings and adults, we made intracellular recordings from them, injected dye, imaged the neurons with a confocal microscope, and traced and measured the neurons (see Methods; statistical comparisons between hatchling and adult neurons are given in Table 2). The anatomy of olfactory PNs in hatchlings resembled scaled-down versions of those in the adult, with spoke-like radiating dendrites ending in glomerular tufts in the AL, an axon projecting to the calyx of the mushroom body and forming a nest-like branching pattern, and a neurite continuing to and terminating with a thicket of branches in the lateral horn (LH). In adults and hatchlings, the numbers of radiating dendritic branches formed by PNs in the AL were about the same (14.6 ± 2.7 in hatchlings, 15.4 ± 2.5 in adults, mean ± SEM, t(5)=0.22, p=0.83), and similarly, the number of glomeruli per PN, defined as fiber bundles at the terminals of PN dendrites, were about the same, too (13.16 ± 0.79 in hatchlings and 14.68 ± 0.70 in adults, t(19)=0.34, p=0.74). Although resembling those in the adult, neurons in the hatchlings were smaller. For example, the lengths of PN axons were smaller in hatchlings, (401.0 ± 3.3 μm in hatchlings and 768.3 ± 28.6 μm in adults, t(5)=12.75, p<<0.001). Notably, though, PNs scaled by brain size; when an individual’s neuron branch length was normalized by the diameter of its antennal lobe (134 ± 5.8 μm in hatchlings and 288.26 ± 9.0 μm in adults), normalized hatchling and adult values were about the same (4.92 ± 0.44 in hatchlings vs 4.78 ± 1.26 in adults, t(5)=0.10, p=0.92). PN branches in the calyx and the LH scaled similarly with age (Table 2A).

**Table 2:** Quantitative comparison of PNs in locust hatchlings and adults. Numbers indicate mean ± SEM. All lengths are in µm.

| **Table 2A** | Hatchling | Adult | T<br>statistic | p-value | Hatchling<br>normalized | Adult<br>normalized | T<br>statistic | p-value |
| --- | --- | --- | --- | --- | --- | --- | --- | --- |
| Total length of PN branches in AL | 659.7±89.5 | 1376.7±353.<br>3 | 1.97 | 0.08 | 4.92 ± 0.44 | 4.78 ± 1.26 | 0.10 | 0.92 |
| Length of PN axon | 401.0±3.3 | 768.3±28.6 | 12.75 | <<0.001 | 2.99 ± 0.13 | 2.67 ± 0.06 | 2.23 | 0.06 |
| Total length of PN branches in MB | 2986.7±76.6 | 6753.1±862.<br>5 | 4.35 | 0.002 | 22.29 ± 0.82 | 23.43 ± 2.56 | 0.42 | 0.68 |
| Total length of PN branches in LH | 693.5±101.7 | 1584±168.1 | 4.53 | 0.002 | 5.18 ± 0.79 | 5.48 ± 0.52 | 0.31 | 0.76 |

| **Table 2B** | Hatchling | Adult | T<br>statistic | p-value |
| --- | --- | --- | --- | --- |
| Diameter of AL | 134 ± 5.8 | 288.26 ± 9.0 | 14.41 | <<0.001 |
| No. of dendritic PN branches in AL | 14.6 ± 2.7 | 15.4 ± 2.5 | 0.22 | 0.83 |
| No. of PN branches in MB calyx | 88.4±2.2 | 93.0±14.6 | 0.31 | 0.76 |
| No. of spines in PN branches in MB | 170.0±10.9 | 206.4±35.5 | 0.98 | 0.36 |
| No. of PN branches in LH | 24.4±5.0 | 27.2±3.2 | 0.47 | 0.65 |
| No. of spines in PN branches in<br>LH | 42.4±8.1 | 56.6±7.7 | 1.27 | 0.24 |
| No. of glomeruli in AL | 13.6±2.2 | 14.6±2.0 | 0.34 | 0.74 |

Quantitative analysis of LNs in hatchlings and adults was challenging because these neurons have diverse and dense morphologies including numerous very fine processes, making them difficult to completely trace and compare with one another. Nevertheless, visual inspection of many filled neurons indicated that hatchling LNs, like PNs, are smaller versions of the adult form. Figure 3 shows representative examples of LN morphologies in hatchlings and adults. Throughout development these neurons typically span the entire AL with dense branches radiating from the center. Summary information for a sampling of LNs that could be traced is provided in Table 3.

**Figure 3:**
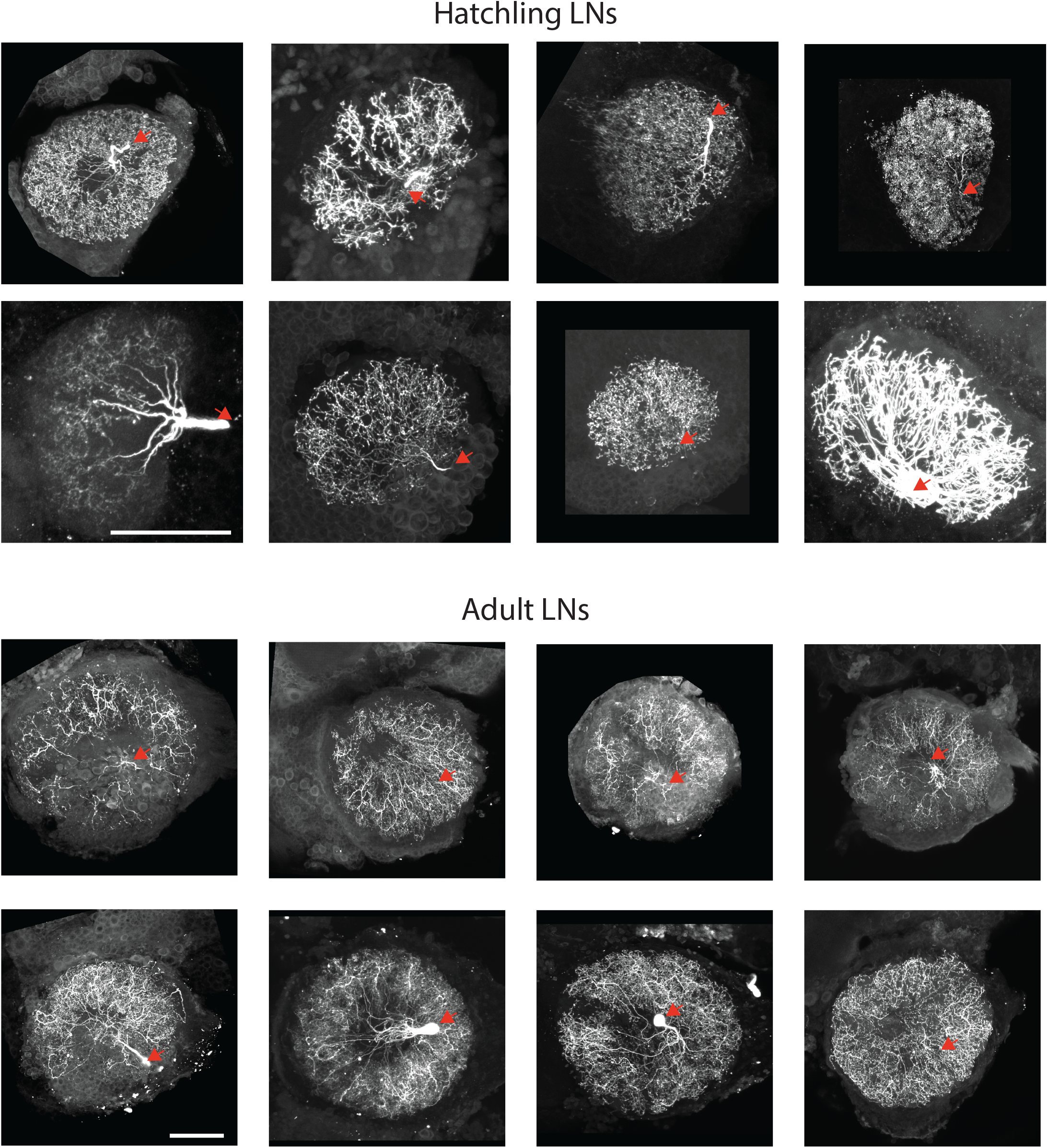
Representative examples of LNs in locust hatchlings (top) and adults (bottom). The images show the maximum intensity projections of the AL containing the dye-filled cells. Red arrow: location of the cell body which is often detached when the patch electrode is removed. Scale bar: 100 μm.

**Table 3:**
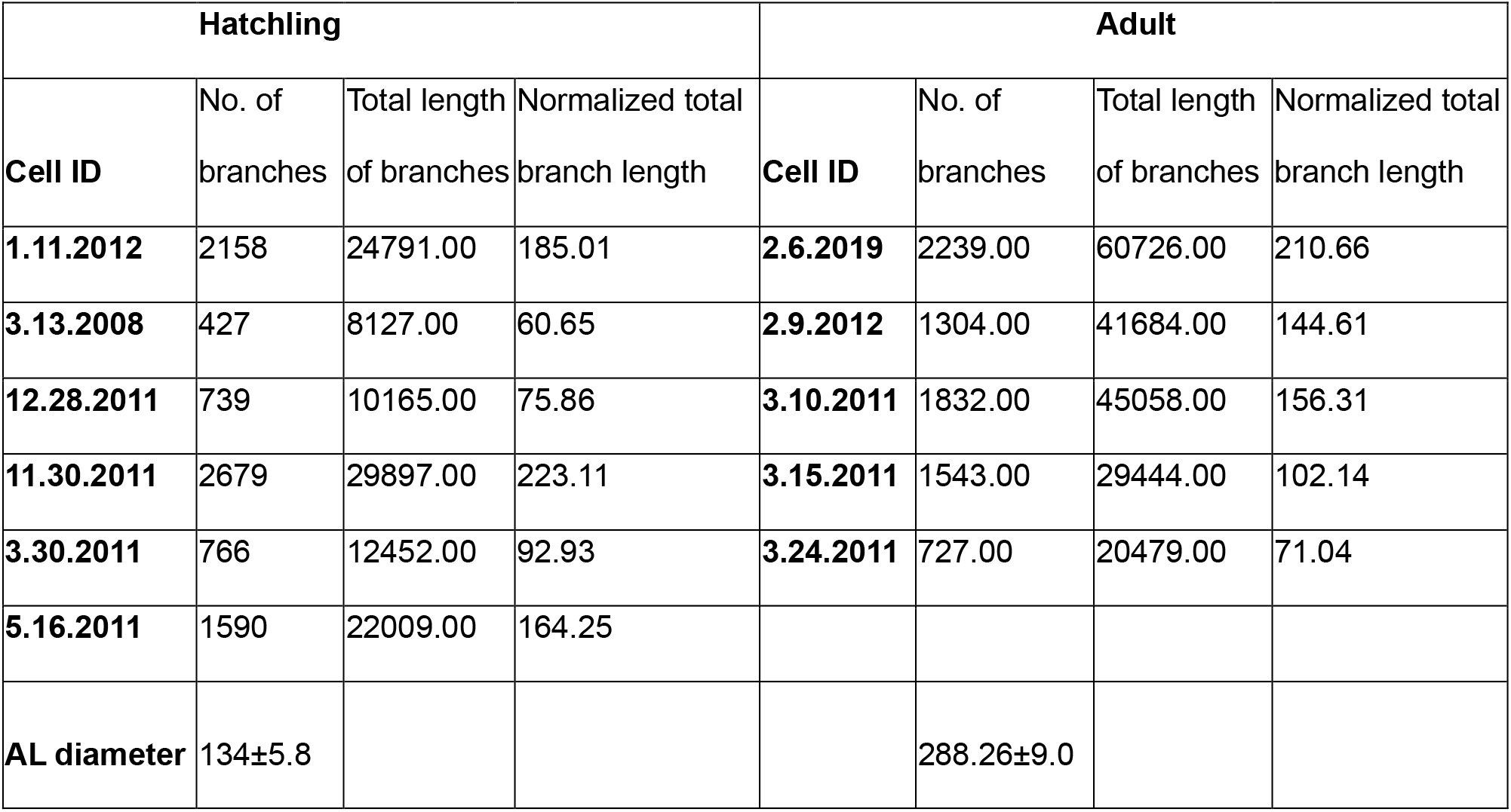
Comparison of the morphologies of traced LNs. All lengths are given in µm. Lengths were normalized by the average diameter of the antennal lobes.

| Hatchling |  |  |  | Adult |  |  |  |
| --- | --- | --- | --- | --- | --- | --- | --- |
| Cell ID | No. of branches | Total length of branches | Normalized total branch length | Cell ID | No. of branches | Total length of branches | Normalized total branch length |
| 1.11.2012 | 2158 | 24791.00 | 185.01 | 2.6.2019 | 2239.00 | 60726.00 | 210.66 |
| 3.13.2008 | 427 | 8127.00 | 60.65 | 2.9.2012 | 1304.00 | 41684.00 | 144.61 |
| 12.28.2011 | 739 | 10165.00 | 75.86 | 3.10.2011 | 1832.00 | 45058.00 | 156.31 |
| 11.30.2011 | 2679 | 29897.00 | 223.11 | 3.15.2011 | 1543.00 | 29444.00 | 102.14 |
| 3.30.2011 | 766 | 12452.00 | 92.93 | 3.24.2011 | 727.00 | 20479.00 | 71.04 |
| 5.16.2011 | 1590 | 22009.00 | 164.25 |  |  |  |  |
| AL diameter | 134±5.8 | | | | 288.26±9.0 | | |

### Hatchling and adult AL neurons respond similarly to olfactory stimuli

We next used patch clamp electrophysiology to systematically compare odor-elicited responses in AL neurons in newly hatched and adult locusts. We first recorded from 15 PNs each in hatchling and adult locusts as they responded to a panel of odors, each puffed into a constant airstream directed at the antennae (see Methods).

Neurons in hatchling and adult locusts responded to odors with comparable assortments of firing patterns including complex sequences of excitation, inhibition, and quiescence. Figure 4A shows examples of these firing patterns as voltage traces obtained by intracellular recording; Figure 4B shows the reliability of these responses with additional examples shown as spike rasters from 15 PNs each in adult and freshly hatched locusts responding to 8 odorants presented 6 times each. Notably, the spontaneous and odor-elicited firing rates of hatchling PNs was significantly lower than that of the adult (Figure 4C); a 2-way ANOVA revealed a significant effect of age (F=35.87, df=1, p << 0.001) with no significant effect of odor or the interaction term, p=0.38 and 0.80 respectively).

**Figure 4:**
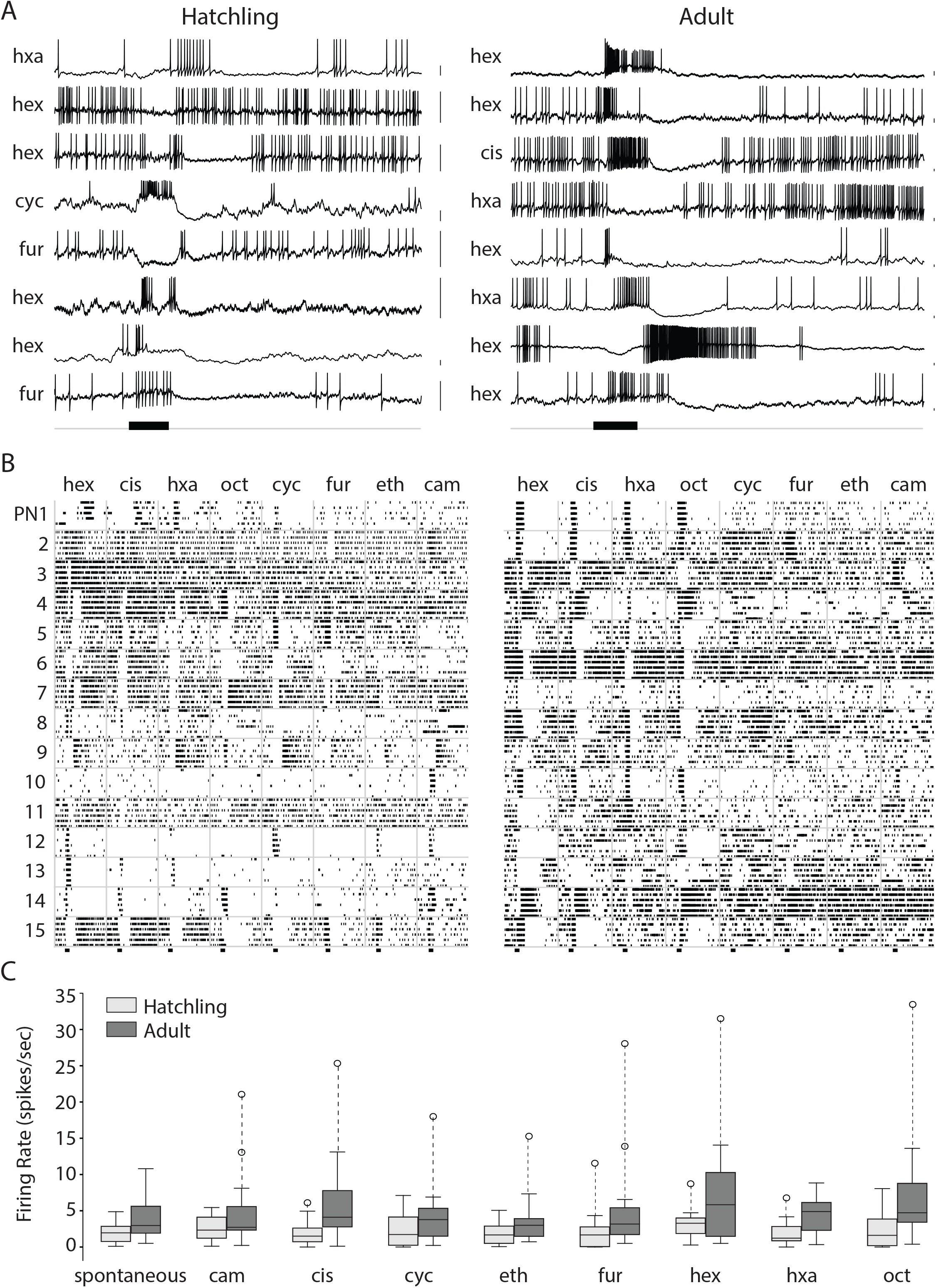
Odors elicit similar patterns of spikes in hatchling and adult PNs, although hatchling PNs responses contain fewer spikes. (a) Representative example membrane potential records from PNs in the AL of both hatchling and adult locusts show combinations of excitation and inhibition that vary with the odor and the cell. Black horizontal bar: 1 sec odor pulse. Odors: hex, hexanol; cis: cis-3-hexen-1-ol; hxa: hexanal; oct: octen-3-ol; cyc: cyclohexanone; fur: furfuryl mercaptan; eth: ethyl mercaptan; cam: camphor. Scale bars: 5 mV. (b) Additional examples: raster plots show spike trains in 15 PNs each in adults and hatchlings tested with 8 different odors. (c) Hatchling PNs generated fewer spikes spontaneously and in response to odors than adult PNs; see text for statistical analysis.

We compared the complexity of odor-elicited spiking patterns in PNs by computing the number of transitions between high and low firing rates within a 4 s time window from odor onset and compared the distribution of the number of transitions across trials (n=414 in adults and 398 in hatchlings; Figure 5; (Raman et al., 2010)). A Kolmogorov-Smirnov test comparing two empirical distributions revealed no significant differences between the two age groups (KS statistic=0.04, p=0.84). This analysis suggests olfactory circuitry in the hatchling brain, though smaller in scale, can generate complex odor-elicited sequences of spiking responses like those observed in adult PNs.

**Figure 5:**
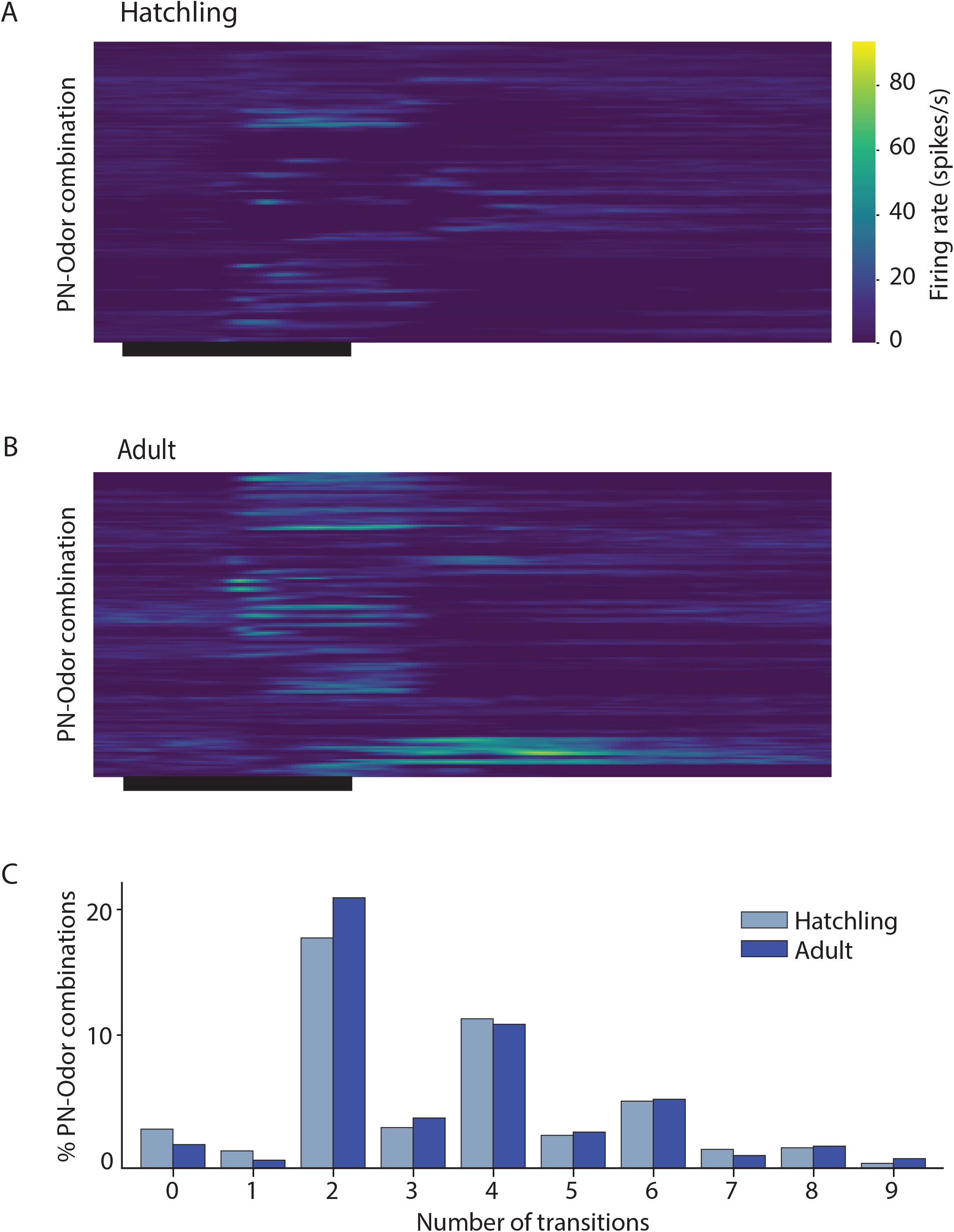
PNs in the hatchling antennal lobe produce complex multiphasic patterns of firing like that in the adult. (a) and (b) Transitions between excitatory and inhibitory epochs in odor-elicited responses of hatchling and adult PNs. Color indicates firing rate in spikes/s; black horizontal bar: 1 sec odor pulse. (c) PNs in adults and in hatchlings did not differ significantly in response complexity as defined by the number of epoch transitions; see text for statistical analysis.

We then made patch clamp recordings from LNs. In locusts, LNs respond to odors with a variety of patterns of depolarization, hyperpolarization, ∼20 Hz membrane potential oscillations, and small calcium spikelets (Laurent & Davidowitz, 1994; MacLeod & Laurent, 1996). The representative examples shown in Figure 6 reveal LNs in newly hatched and in adult locusts display a similar array of response types to puffs of odorants.

**Figure 6:**
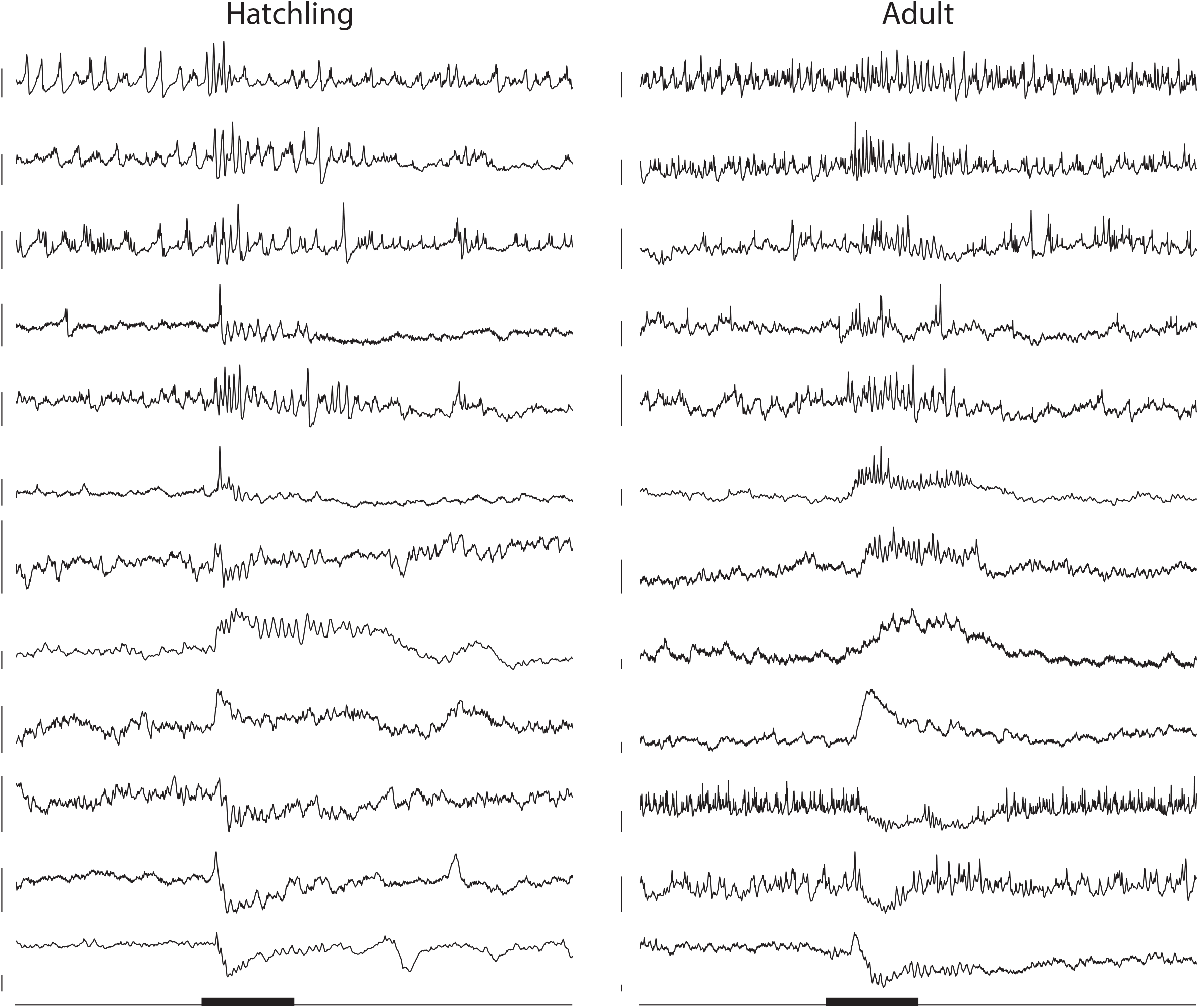
LNs in hatchlings and adults respond similarly to odors. Representative examples of membrane potentials of LNs in hatchling and adult locusts responding to 1 sec odor pulses (black bar). Each trace is from a different LN. Locust LNs do not produce action potentials, but rather calcium spikes, the rate of which may change upon odor presentation. LNs membrane potential can also oscillate, depolarize, hyperpolarize, or undergo a mixture of these. LNs in animals of both age groups show similar responses. Scale bars: 10 mV.

### Odor-evoked LFP oscillations are slower in hatchlings

In adult locusts, odor presentations are known to elicit the synchronized and oscillatory spiking of populations of PNs centered around 20 Hz, a phenomenon that can be assessed by local field potential (LFP) recordings made by extracellular electrodes in the AL and the MB (Laurent & Naraghi, 1994; Stopfer & Laurent, 1999). To compare the coordination of AL neurons at different ages, we recorded odor elicited LFPs in the MBs of freshly hatched, 10 days old, 20 day old, and 35 day old animals, corresponding to first, third, and fifth instar larval and adult locusts. LFP oscillations were evident in the hatchlings, indicating that even at this early developmental stage the reciprocal circuitry between LNs and PNs in the AL necessary for the generation of oscillations was already present and appropriately tuned (Bazhenov et al., 2001). However, we found, over the course of development, that oscillations changed in three ways: the frequency and duration of the oscillations gradually increased; and the delay between odor delivery and oscillation onset increased (Figure 7).

**Figure 7:**
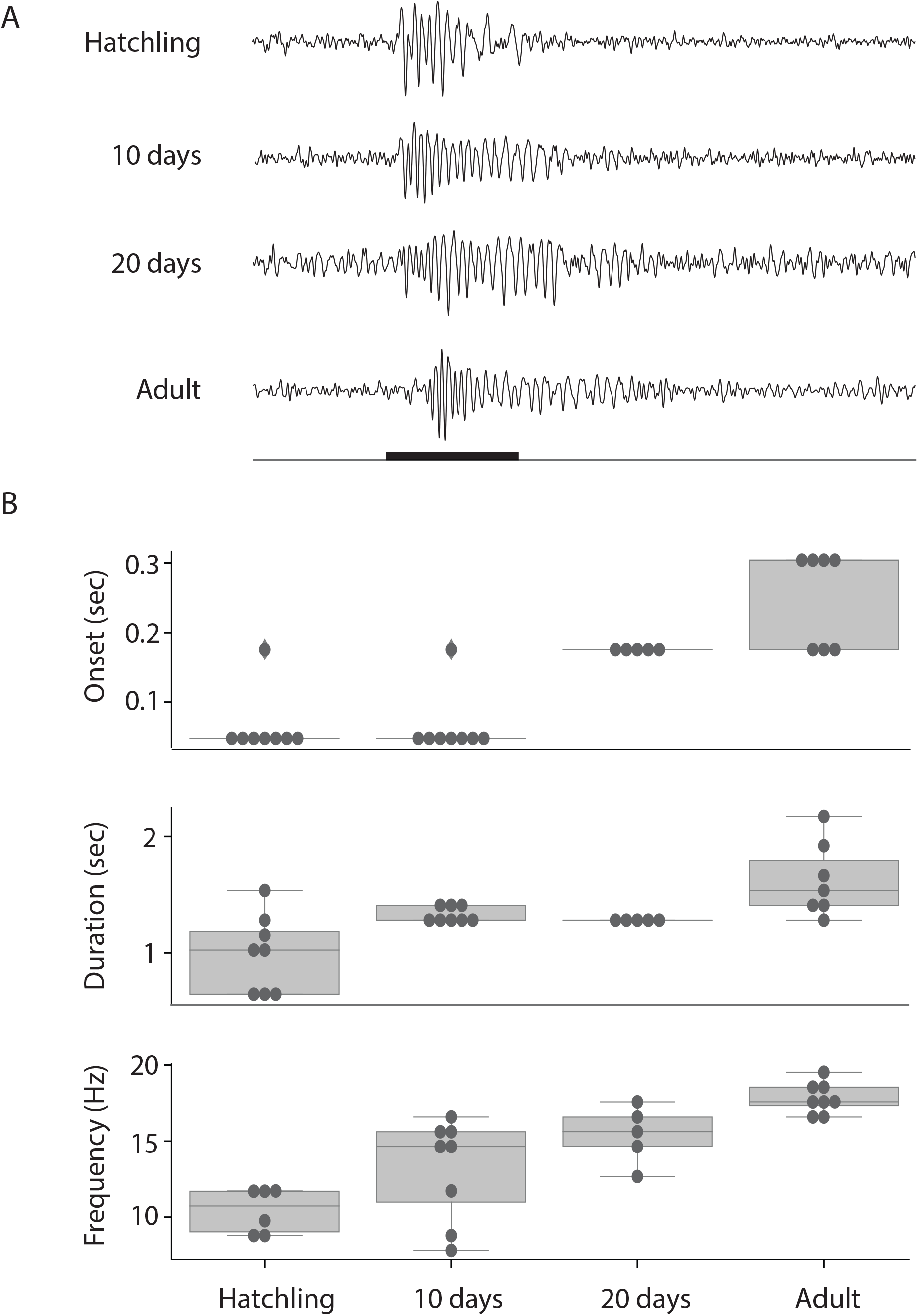
In response to odor puffs, hatchlings generate lower-frequency LFP oscillations than adults. (a) Representative LFP oscillations recorded in the MBs of hatchling, 10 day old, 20 day old, and adult locusts; Black horizontal bar: 1 sec pulse of 100% hexanol. (b) properties of LFP oscillation across these age groups. Boxes cover first to third quartile, interior line: median, whiskers: 1.5 x inter-quartile range, circles: individual datapoints, and diamonds: outliers.

### Antenna length correlates with LFP oscillation frequency

What mechanism underlies these age-dependent shifts in odor-elicited oscillatory activity? Earlier work established that, in the AL, oscillation frequency is determined by the intensity of odor-elicited excitation from the afferent layer driving the oscillator circuitry (Ito et al., 2009). Locusts add OSN-containing segments to their antenna as they develop, beginning with approximately 8,000 OSNs spread over 11-13 segments in hatchlings, and increasing to over 80,000 OSNs spread over 24-26 segments in adults (Kuitert & Connin, 1952; Greenwood & Chapman, 1984; Chapman & Greenwood, 1986; Capinera, 1993; Ochieng et al., 1998; Chapman, 2002). The evident relationship between the developmental increase in the number of OSNs and the developmental increase in oscillation frequency led us to hypothesize that oscillation properties might be determined by the number of OSNs driving the oscillator circuit. To test this, in adult locusts, we trimmed successive antennal segments while recording odor-elicited LFPs in the MB ipsilateral to the antenna. As a within-animal control, we left the contralateral antenna intact while also recording odor-elicited LFPs from that side of the brain (Figure 8a). We found that, as segments were removed, the LFP oscillation frequency measured in the ipsilateral MB significantly decreased, whereas the frequency remained unchanged on the intact side (Figure 8b, c). A linear regression (ordinary least squares fit) of the number of cuts to the peak LFP frequency during odor presentation showed significant negative correlation with the number of cuts for ipsilateral side (coefficient=-.52±0.11, R^2^=0.258, p << 0.01), but not for the contralateral side (coefficient= 0.06±0.079, R^2^=0.023, p=0.445). To confirm that this reduction of LFP frequency was due to the progressive reduction in number of OSNs and not by gradual systemic degradation following injury, we also performed quicker experiments where we directly cut the antenna after the 7^th^ segment (leaving 19 segments intact), and then after the 10^th^ segment (leaving 16 segments intact, Supplementary Figure 1); this approach yielded the same result.

**Figure 8:**
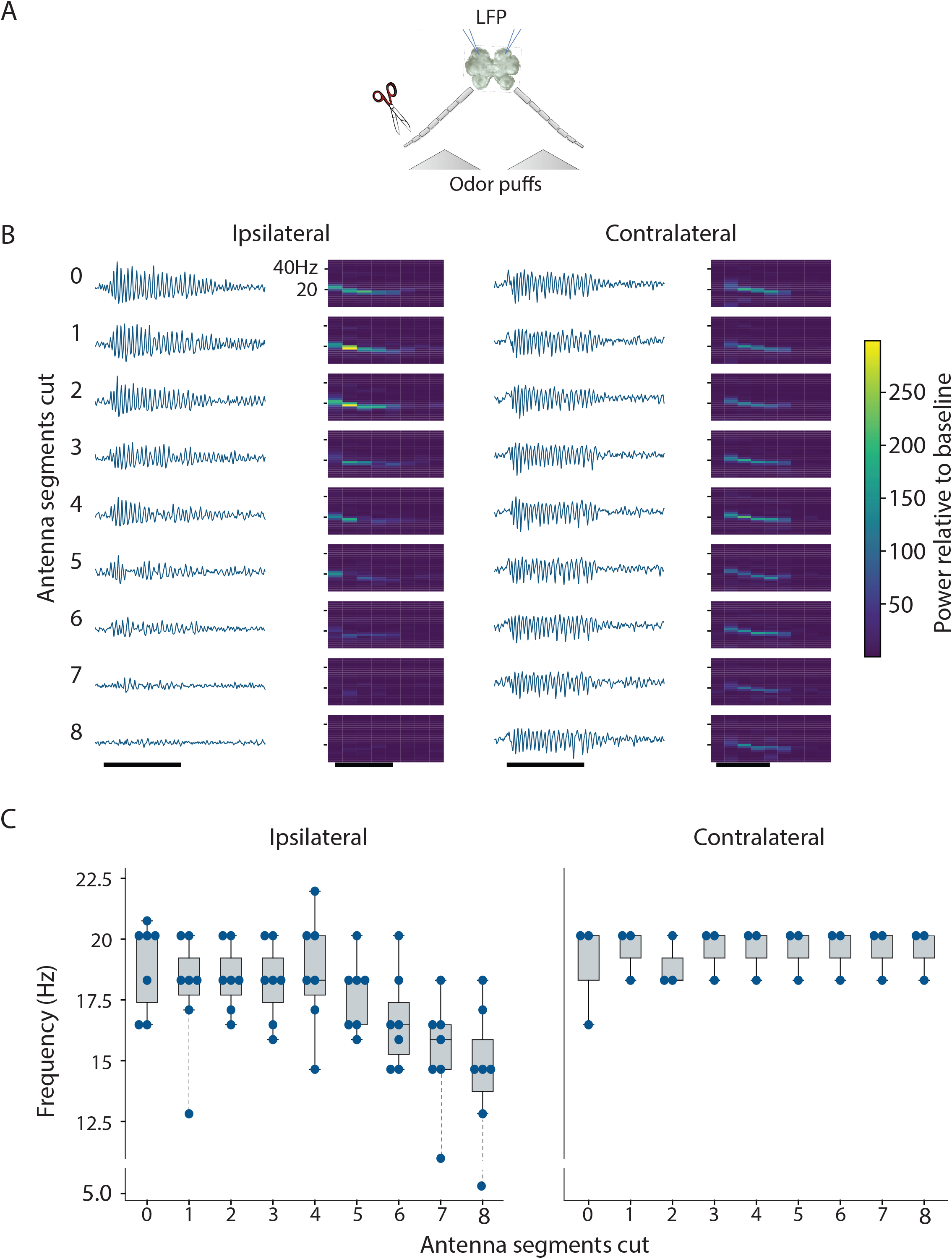
LFP oscillation frequency correlates with the number of ORN-containing antennal segments; removing segments from adults reduced oscillation frequency. (a) Experiment scheme for trimming antennal segments from one side of an adult animal while recording LFPs from the ipsilateral MB; the contralateral antenna and MB served as a within-animal control. (b) Odor-evoked LFPs and corresponding spectrogram in the MB ipsilateral (left) to the trimmed antenna (left half) and the same in the contralateral AL (right) of a single animal. Black horizontal bars: 1 sec odor presentation. (c) As antennal segments were trimmed, LFP frequency in the ipsilateral MB decreased (left), but the frequency in the contralateral MB remained unchanged (right) (n=7 antennae in 4 animals). Blue dots: individual data points; boxes cover first to third quartile, interior line: median, whiskers: 1.5 x inter-quartile range; dotted line: outliers. Each data point shows the frequency with the peak power sampled 2-3 sec in the averaged spectrogram of trials 3 to 10. First two trials were excluded because oscillations usually develop after 2 odor presentations. Odorant: hexanol.

## Discussion

Locusts have served as useful models to understand how olfactory stimuli are encoded by neural circuits (Laurent & Davidowitz, 1994; Laurent, 1999; Laurent et al., 2001; MacLeod & Laurent, 1996; Mazor & Laurent, 2005; Perez-Orive et al., 2002; Stopfer et al., 1997; Stopfer & Laurent, 1999; Stopfer et al., 2003; Uchida et al., 2014). Here we investigated the development of peripheral stages of the olfactory pathway by comparing naïve, freshly hatched, and adult locusts. One might hypothesize that developmental growth or experience with the olfactory environment would substantially alter the structures and functions of the locust’s olfactory circuitry. However, at the level of confocal microscopy, hatchling olfactory neurons resembled those of the adult, just smaller, scaled by overall brain size. EAG recordings from hatchling and adult antennae revealed OSNs responded with similar sensitivities to a panel of odors across development. A larger odorant panel including pheromones, and recordings from individual OSNs might reveal more specific experience-or age-dependent changes in tuning.

Earlier work has shown that in hemimetabolous insects like locusts, the main structures of the brain, including the antennal lobes and their full complement of glomeruli, are well formed upon hatching (Panov, 1961; Prillinger, 1981; Petersen et al., 1982; Chambille & Pierre Rospars, 1985; Kutsch & Hemmer, 1994), although more subtle changes, such as synaptic rearrangement, have been documented throughout development (Chiba et al., 1988). In an earlier study of locust species *Schistocerca gregaria*, Anton and colleagues (Anton et al., 2002) found that, as the animals matured, the fraction of PNs responding to aggregation pheromone components increased, and the overall response profiles of PNs became less selective. In juvenile locusts these authors also observed that PNs responded to odors with patterns including inhibition and excitation. Our analysis of the temporal dynamics of these responses revealed that olfactory neurons in hatchlings responded to odors very much like those of the adults: PNs fired spikes in similarly complex sequences of excitation, quiescence, and inhibition and coordinated in oscillatory synchrony, and, across development, LNs, which do not generate sodium action potentials in locusts, showed very similar patterns of calcium spikelets atop excitatory and inhibitory membrane fluctuations.

Notably, though, PNs in hatchlings showed less spontaneous and odor-elicited spiking than those in adults, and odor-elicited oscillations were slower in frequency in the hatchling. A previous study (Ito et al., 2009) demonstrated that the olfactory sensory neurons (OSNs) in the moth *Manduca sexta* underwent sensory adaptation when exposed to lengthy olfactory stimuli. Adaptation in OSNs led to a reduction in the strength of their response to odors, and a corresponding decrease in the frequency of LFPs. (This downward frequency shift can also be observed here in spectrogram traces in Figure 8B). A computational model incorporating OSN adaptation showed this intensity reduction slowed the LFP frequency by decreasing drive to the oscillator circuitry in the AL (Ito et al., 2009). Here, we hypothesized that LFP oscillations are slower in hatchlings than adults because hatchlings have fewer OSNs, collectively contributing less excitatory drive to the AL oscillatory circuit. To test this idea, we sought to return the adult antenna to the hatchling state by removing successive segments, and thus OSNs, from the antenna. Consistent with our hypothesis, we found LFP oscillation frequency decreased as segments of the adult antenna were removed. We speculate the relatively low excitatory drive from the hatchling antenna may also underlie the lower levels of spontaneous and odor-elicited activity observed downstream in hatchling PNs (figure 4C), as spontaneous activity in PNs has previously been shown to arise almost entirely from OSN spiking (Joseph et al., 2012).

In postembryonic rats, odor elicited oscillations in the olfactory cortex (analogous to the insect mushroom bodies; Barnum & Hong, 2022), undergo a frequency shift during development similar to what we characterize here in locusts. In rats, oscillations shift from 12-15 Hz peak in P0-P12 animals, to ∼25 Hz by P20 (Zhang et al., 2021). However, the frequency transitions in locusts and rats are caused by very different processes: in locusts, the transition is due to an increase in the number of OSNs, while in rats it reflects changes in centrifugal input from the cortex to the olfactory bulb (Zhang et al., 2021).

In many hemimetabolous insect species, although the central brain is fully formed at hatching, the number of peripheral sensory neurons increases through development; visual (Anderson, 1978; Stark & Mote, 1981), auditory (Ball & Young, 1974; Young & Ball, 1974; Petersen et al., 1982), wind-sensitive (Chiba et al., 1992; Kutsch & Hemmer, 1994), and olfactory sensory neurons (Chapman & Greenwood, 1986) have all been observed to increase in quantity throughout postembryonic life. As additional sensory neurons are added in these systems, the combined strength of their outputs onto follower neurons increases (Petersen et al., 1982; Bucher & Pflüger, 2000). This matches our observations in locusts, where increased odor elicited EAG amplitude occurs together with increased spontaneous and evoked firing in PNs. The increased sensory drive onto oscillator circuits in the AL also led to increased oscillatory frequency measured in the LFP. This phenomenon may be analogous to burst frequency in motor circuits organizing wing beats in the cricket, which increases through development as sensory input increases (Bentley & Hoy, 1970).

Notably, as we observed in the elaborate firing sequences of PNs and the oscillatory synchronization of AL neurons, the response properties of olfactory neurons remain consistent throughout development, despite dramatic changes in both the intensity of sensory drive (Chiba et al., 1992), and the size of the follower neurons (Anderson, 1978; Stark & Mote, 1981). In cockroaches, growing neurons adjust the lengths and diameters of their processes to maintain consistent electrical properties (Hochner & Spira, 1987), whereas crustacean neurons appear to achieve functional stability throughout development by adjusting membrane conductances (Liu et al., 1998). It would be interesting to determine how the locust antennal lobe generates consistent output despite dramatic developmental changes in size and input.

In other species, the olfactory system has shown only a limited ability to change in response to early life exposure to odorants, including maintaining memories formed during development (Blackiston et al., 2008; Arenas et al., 2009; Iyengar et al., 2010). However, it is not clear that such plasticity, shown to occur under laboratory conditions, is functionally relevant to animals under more natural conditions (Gugel et al., 2023).

In summary, our results show that locusts hatch with a well-developed olfactory neural circuit – one with neurons resembling and functioning like those of the adult. Despite their small size and their lack of prior experience with odorants, OSNs in hatchlings respond with similar tunings to odorants, and second order neurons display complex temporal patterns of activity and organize into oscillating ensembles in response to olfactory stimulation. These results are consistent with our earlier finding that newly hatched locusts are innately attracted to the odors of food sources (Ray et al., 2023). Fully assembled olfactory systems help enable hatchling locusts and other hemimetabolous insects to interact effectively with their environments.

## Author contribution

KS and conducted experiments and analyzed data, NG designed the project, conducted experiments and analyzed data, SR analyzed data and wrote the manuscript, ZA conducted experiments, and MS designed and supervised the project and wrote the manuscript.

## Acknowledgements

We thank Vincent Schram and Lynn Holtzclaw of NICHD Microscopy Core for their help with imaging, Bruce Pritchard, George Dold, and members of NIMH Section on Instrumentation for help with experimental setup, and members of the Stopfer Lab for constructive comments and suggestions.

---

**Supplementary Figure 1:**
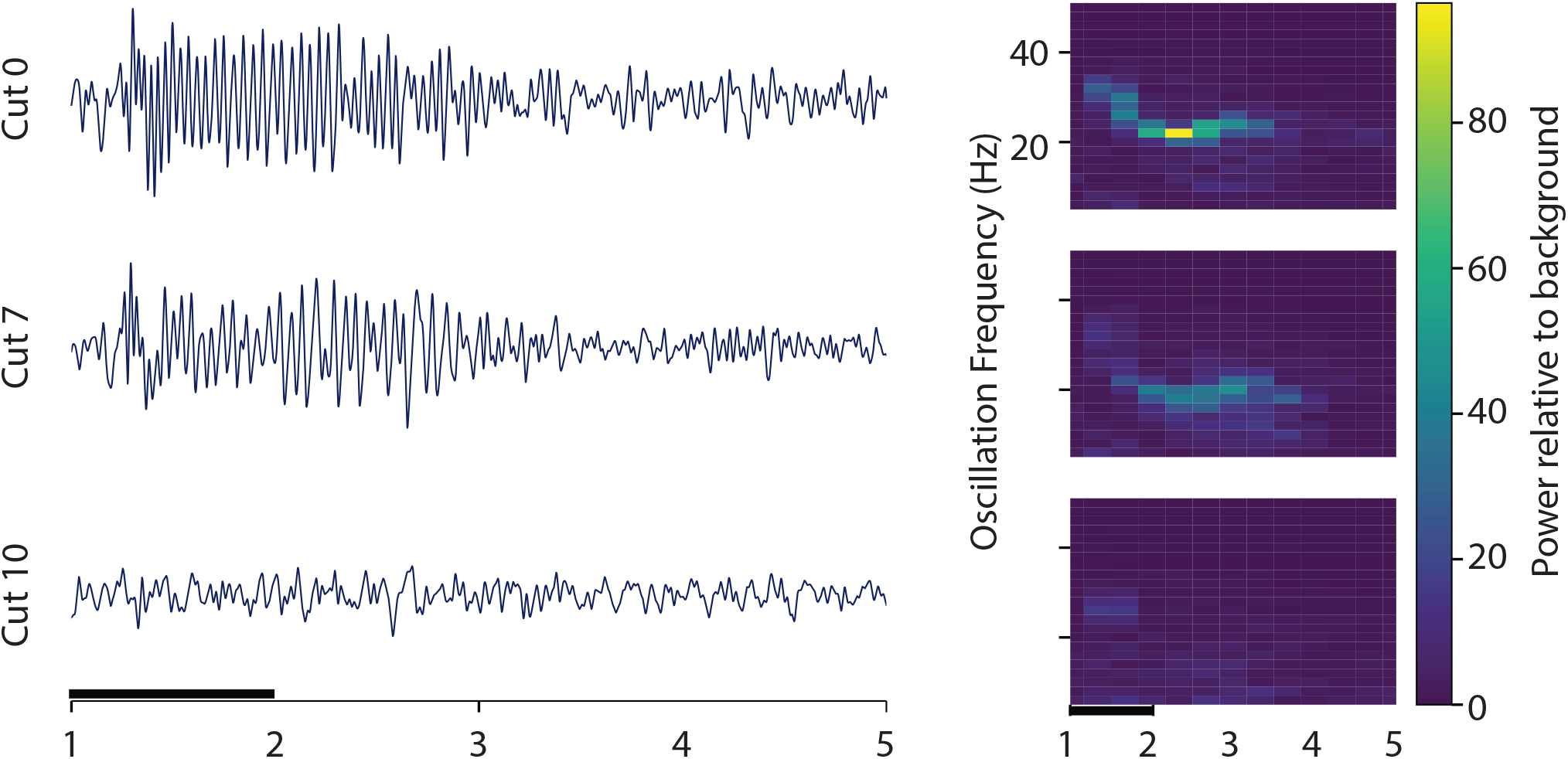
Left: LFP response when the antenna of an adult locust is cut after segments 7 and then after segment 10. Right: Spectrogram of the LFP response shows decreasing oscillation frequency as antennal segments are removed. Black horizontal bar: 1 sec odor pulse.

